# Evaluation of the DBA/2J mouse as a potential background strain for genetic models of cardiomyopathy

**DOI:** 10.1101/2022.05.16.492163

**Authors:** Cora C. Hart, Young il Lee, David W. Hammers, H. Lee Sweeney

**Author notes:** Equal contributions by these authors (David W. Hammers, H. Lee Sweeney). Corresponding author, **Correspondence:** H. Lee Sweeney, 1200 Newell Dr. ARB R5-216, Gainesville, FL 32610-0267.

## Abstract

The potential use of the D2.*mdx* mouse (the *mdx* mutation on the DBA/2J genetic background) as a preclinical model of the cardiac aspects of Duchenne muscular dystrophy (DMD) has been criticized based on speculation that the DBA/2J genetic background displays an inherent hypertrophic cardiomyopathy phenotype. Accordingly, the goal of the current study was to further examine the cardiac status of this mouse strain over a 12-month period. DBA/2J mice have been scrutinized for the presence of cardiac lesions, however, in the current study we find that DBA/2J mice contain equivalent amounts of left ventricular collagen as healthy canine and human samples. In a longitudinal echocardiography study, neither sedentary or exercised DBA/2J mice demonstrated left ventricular wall thickening or cardiac functional deficits. In summary, we find no evidence of hypertrophic cardiomyopathy, or any other cardiac pathology, and thus propose that it is an appropriate background strain for genetic modeling of cardiac diseases, including the cardiomyopathy associated with DMD.

## INTRODUCTION

Effective preclinical testing of potential therapeutics to treat human diseases requires the use of a model that accurately represents a disease progression and severity that is homologous to the human condition. For example, Duchenne muscular dystrophy (DMD) is the most frequently inherited pediatric neuromuscular disease, however, there remains limited treatment options for this patient population despite almost three decades of preclinical and clinical research. In fact, 16 compounds have been discontinued during clinical development for DMD and this has largely been attributed to the lack of an animal model that is representative of the human disease [1].

Indeed, the most widely used preclinical model of DMD is the *mdx* mouse on C57-based genetic backgrounds [2, 3], despite their relatively mild phenotype in the limb muscles and little discernable cardiac disease until they reach old age. The increased cardiac pathology of older *mdx* mice is likely due to aging-related factors combined with dystrophin deficiency. Such age-associated factors include the diastolic dysfunction that is evident in aged wild-type mice [4]. This has led to the utilization of exogenous means, such as β-adrenergic stimulation, to worsen phenotype beyond the natural disease progression, differentiate wild-type and dystrophic cardiac function, and/or identify potential therapeutic effects [5–8]. Additional genetic manipulations, such as ablation of utrophin [9] or telomerase [10, 11], have been used to worsen skeletal and cardiac muscle pathologies and decrease life expectancy of C57-based *mdx* mice. However, these manipulations do not represent genetic homologs of DMD and may mask potential therapeutic effects and/or limitations. For example, we have noted utrophin-mediated cardiac benefits following treatment with a PDE5 inhibitor [5] that would not be seen in a utrophin-deficient model. Therefore, in order to conduct translatable preclinical studies, an animal model that recapitulates both the skeletal and cardiac muscle disease progression of DMD is needed.

Recently, the D2.*mdx* mouse, consisting of the *mdx* mutation on the DBA/2J genetic background, has emerged as a severe mouse model of DMD that exhibits many features of the skeletal muscle pathology seen in human patients, including regenerative defects, progressive muscle fibrosis, muscle wasting, and reduced life-expectancy [12–16]. However, the utility of this mouse line as a suitable model to study the cardiomyopathy associated with DMD has been questioned, particularly over concerns that the DBA/2J genetic background displays inherent pathologies such as cardiac fibrosis [17, 18] and a hypertrophic cardiomyopathy (HCM) [19]. In order to perform rigorous preclinical investigations, isogenic background strains devoid of any confounding pathology are necessary. Therefore, we sought out to carefully examine the cardiac status of DBA2/J (D2.WT) mice. As we detail below, we find no evidence of pre-existing cardiac pathology and thus assert that this genetic background is a viable model to study cardiomyopathy-causing genetic diseases, such as in DMD.

## RESULTS

A feature of D2.WT hearts that has been interpreted as pathological, and possibly an indication of inherent HCM development, is the aberrant extracellular matrix (ECM) deposition [17, 19]. The DBA/2J background contains a polymorphism in latent TGFβ binding protein 4 (LTBP4) that increases the susceptibility of TGFβ activation over that of mice from C57-based backgrounds [20]. While the difference between the C57 and DBA/2J LTBP4 sequence has been a point of concern, human LTBP4 more closely resembles the DBA/2J variant than C57 LTBP4 [21]. We examined whether the supposed excessive cardiac ECM deposition downstream of heightened TGFβ activity in DBA/2J background is pathological. As shown in the picrosirius red stained image of **Figure 1a-b**, D2.WT hearts do exhibit a dense epicardial patch covering the right ventricle (RV), whereas other regions of the heart, particularly the left ventricle (LV) and interventricular septum (**Figure 1c-d**), contain observable, but not pathological, ECM/connective tissue surrounding blood vessels and cardiomyocyte bundles. Previous investigations into the RV epicardial patch of D2.WT mice concluded that the presence of this structure does not impair cardiac function or affect mouse life span [18]. Similar epicardial regions and ventricular organization patterns as those observed in D2.WT mice can be found in heart samples from normal (cardiomyopathy-free) canines (**Figure 1e**). Furthermore, regions of B10.WT hearts exhibiting disorganization and perivascular ECM are also observed (**Figure 1f**), demonstrating these features are not unique to the DBA/2J genetic background.

**Figure 1.**
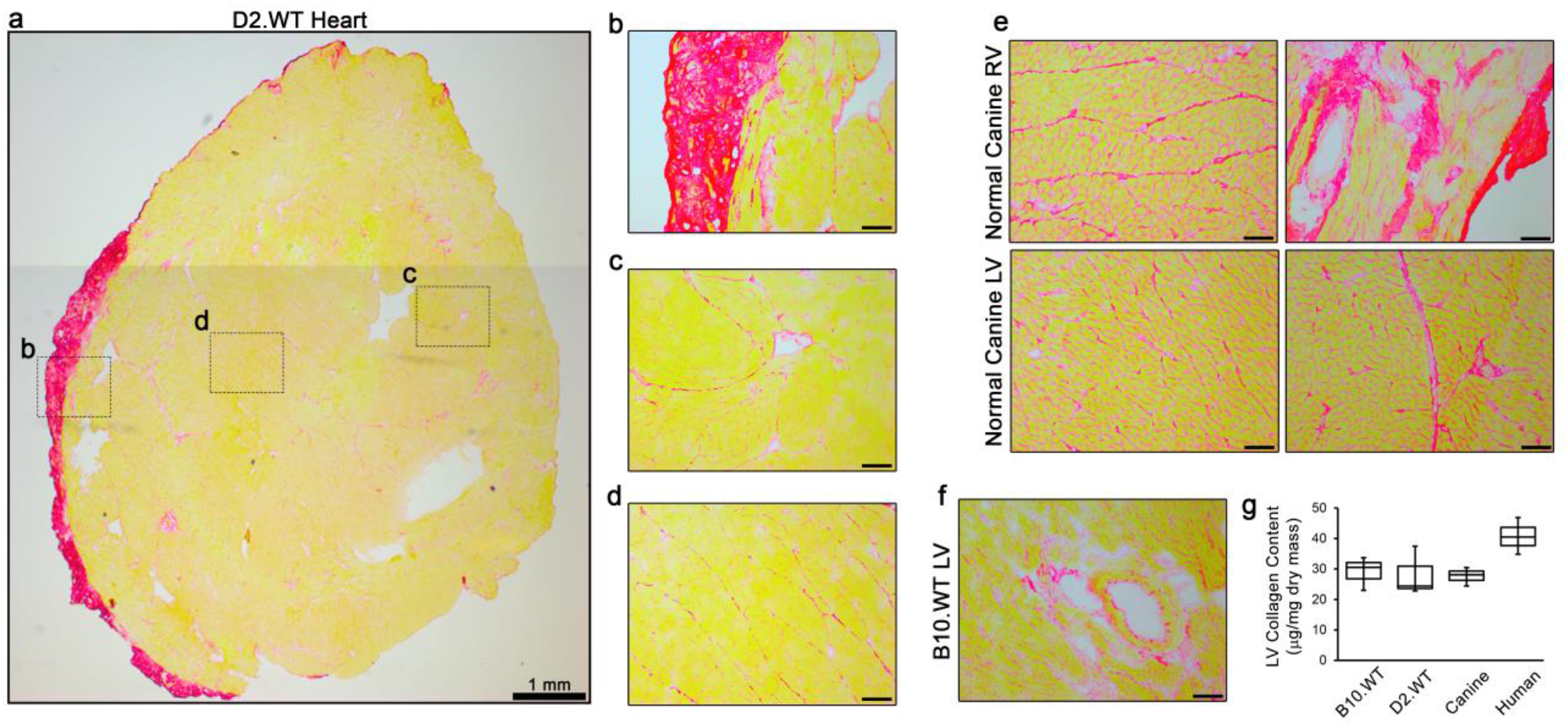
Comparative analysis of left ventricular collagen content. (**a**) Representative picrosirius red staining of a 12 mo D2.WT heart cross-section reveals (**b**) robust collagen staining at the epicardium of the right ventricle (RV) and non-pathological connective tissue organization within the (**c**) left ventricle (LV) free-wall and (**d**) interventricular septum, particularly in areas containing major blood vessels. (**e**) Images of RV and LV samples from 12 mo normal canine hearts reveal similar features as the D2.WT hearts, including regions of dense collagen content of the RV free-wall and perivascular connective tissue. (**f**) The vessel-associated collagen detection by picrosirius red staining is also featured in the hearts of 12 mo B10.WT mice. (**g**) Biochemical analysis of LV collagen content from 12 mo B10.WT, 12 mo D2.WT, 6-12 mo normal canine, and 20-34 yo human hearts (n = 3). Data were analyzed using ANOVA. No significant differences between groups were found. Scale bars indicate 100 μm, unless otherwise noted.

To determine how LV collagen content compares between different mouse strains and species, biochemical quantification of collagen content was performed for LV samples from C57BL/10 (B10.WT) and D2.WT mice (12 months of age), as well as normal canines (6-12 months of age) and humans having no history of cardiomyopathy (20-34 years of age). Interestingly, LV collagen content (normalized to dry tissue mass) was nearly identical between B10.WT, D2.WT, and canine samples, with a slight (non-significant) increase found in human samples (**Figure 1g**). This indicates that excessive and pathological levels of LV fibrosis is not a feature of D2.WT hearts.

To specifically investigate the validity of HCM development in this mouse line, we utilized *in vivo* echocardiography to directly assess cardiac structure and function. HCM and exercise have long been thought of as incompatible [22], therefore, a cohort of D2.WT mice with *ad libitum* access to a running wheel were included in this study, as previously described [23]. Running wheels were added to the cages at 5.5 weeks of age, and mice ran an average of ~5km per day. A longitudinal study was performed in which sedentary (normal ambulation) and wheel-running mice underwent echocardiography bi-monthly from 4 to 12 months of age.

Because a true HCM phenotype entails the progressive thickening of the left ventricle (LV) and enlargement of cardiomyocytes [24], we sought to determine if D2.WT mice of either activity level exhibit such clinical features of HCM. During the first 12 months of life, we did not observe LV wall thickening or increased LV mass in D2.WT mice, even with increased physical activity (**Figure 2a-b**). Wheel running induced a physiological increase in LV chamber size during diastole at 6 months of age (LV End Diastolic Volume, EDV; **Figure 2c**) that normalized at the 8-month timepoint. Sedentary D2.WT mice had a slight increase in LV EDV from 6 to 10 months of age (**Figure 2c**), contrary to what would be expected with a HCM phenotype. In agreement with these findings, wheel running resulted in increased cardiac reserve at 6 months of age as demonstrated by an increase in end systolic volume (ESV,**Figure 2d**) and a slight decrease in ejection fraction (EF,**Figure 2e**). Both ESV and EF normalized by the 8-month timepoint and remained within normal values throughout the remainder of the study. Likewise, D2.WT mice of both activity levels maintained their stroke volume (SV,**Figure 2f**) throughout the study. Diastolic function was also assessed by means of color and pulsed-wave doppler of the blood flow through the mitral valve. The myocardial performance index (MPI) is the ratio of ventricular isovolumetric and ejection time and is an index of global ventricular function. An increase in this ratio is indicative of dysfunction and was not observed within the 12-month time frame of this study (**Figure 2g**). Furthermore, there was no change in the LV early/late mitral valve inflow ratio (E/A, **Figure 2h**). In summary, we did not observe altered cardiac function suggestive of HCM in sedentary or wheel-running cohorts of D2.WT mice (**Figure 2a-h** and **Table 1**). As shown in **Figure 2i**, D2.WT mice do have a larger heart to body-size ratio, as compared to wild-type C57BL/10 (B10.WT) mice. However, the larger heart masses do not progress disproportionately with age, from 4 to 12 months of age, compared to those of B10.WT mice. Analysis of cardiomyocyte size revealed the same pattern (**Figure 2j**). This is likely due to heightened TGFβ signaling, a positive regulator of cardiomyocyte size [25], in the DBA/2J genetic background [20]. These findings further indicate that spontaneous development of HCM is not a feature of D2.WT mice.

**Figure 2.**
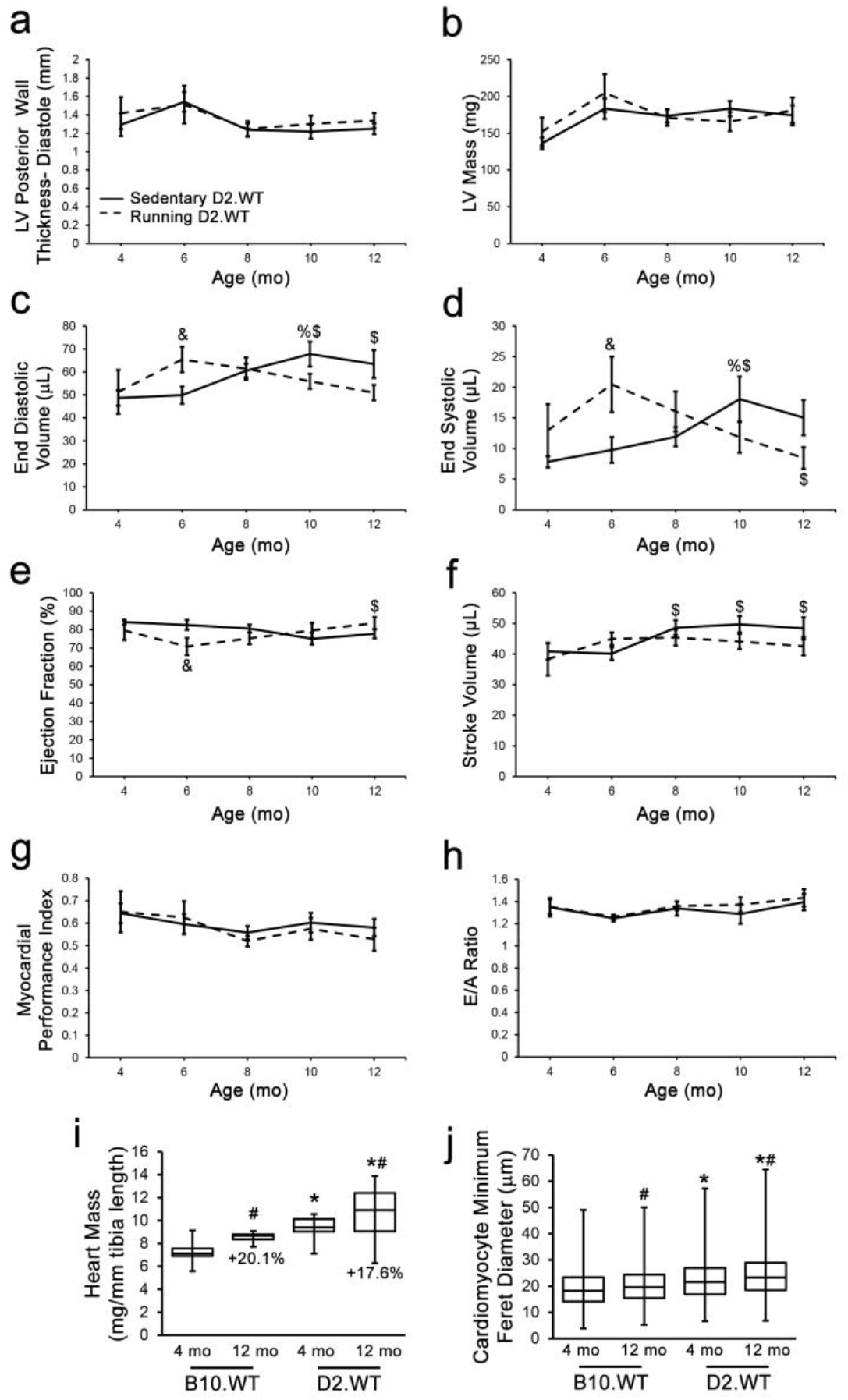
The DBA/2J genetic background does not exhibit hypertrophic cardiomyopathy. Male DBA/2J (D2.WT) mice of sedentary and volitional wheel-running cohorts were followed longitudinally (n = 7-10) between the ages of 4 to 12 months using echocardiography to determine if this mouse strain exhibits a progressive cardiomyopathy resembling hypertrophic cardiomyopathy. Cardiac functional parameters of (**a**) diastolic left ventricle (LV) posterior wall thickness, (**b**) LV mass, (**c**) end diastolic volume, (**d**) end systolic volume, (**e**) ejection fraction, (**f**) stroke volume, (**g**) myocardial performance index, and (**h**) mitral valve E/A ratio from these mouse cohorts are displayed and do not suggest a hypertrophic cardiomyopathy develops in this mouse strain during the ages investigated in this study. A comparison of (**i**) heart mass (normalized to tibia length; n = 6-18) and (**j**) cardiomyocyte size (n = 1066-3919 cardiomyocytes) was performed between 4- and 12-month-old C57BL/10 (B10.WT) and D2.WT mice to identify how heart size progresses between the two genetic backgrounds. (**a-h**) Data were analyzed using two-factor repeated measures ANOVA (Tukey post-hoc tests; α = 0.05); %p < 0.05 vs. group-matched 4 mo values; $p < 0.05 vs. group-matched 6 mo values; & p < 0.05 vs. age-matched sedentary values. (**i-j**) Data were analyzed using two-factor ANOVA (Tukey post-hoc tests; α = 0.05); *p < 0.05 vs. age-matched B10.WT values; #p < 0.05 vs. strain-matched 4 mo values.

**Table 1.**
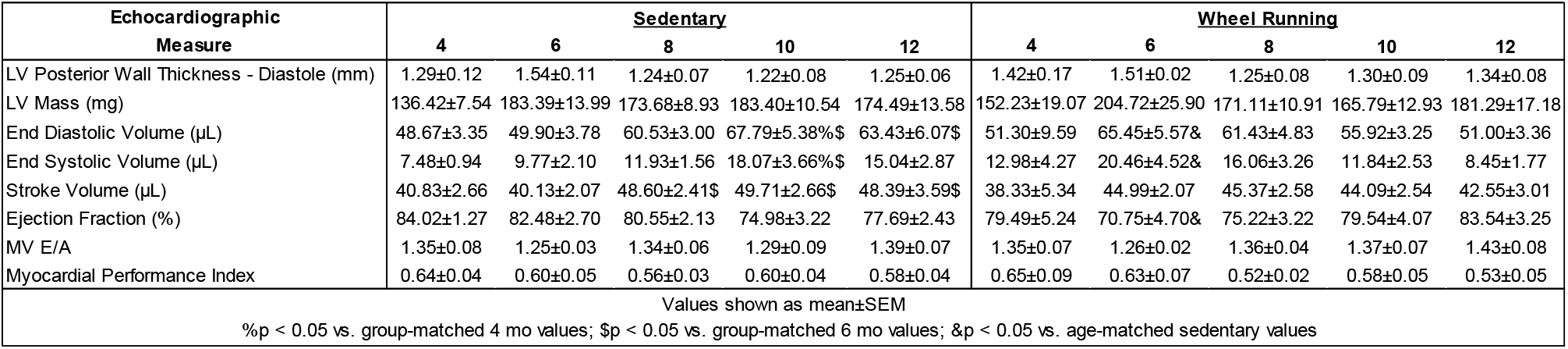
Wild-type DBA/2J Echocardiography.

## DISCUSSION

The data regarding the cardiac status of the DBA/2J mouse have been contradictory [19]. Evidence for the manifestation of HCM in D2.WT hearts reported by Zhao *et al.* [19] was shown at a single time point (4 months of age) and was considered hypertrophic only in comparison with the hearts of C57BL/6 mice. While we also find that the heart mass and cardiomyocyte size of D2.WT mice are indeed larger than those of age-matched B10.WT mice (**Figure 2i-j**), this does not appear to be pathological in nature. It is likely that the heightened TGFβ signaling in D2.WT mice during development and normal postnatal growth results in larger cardiomyocytes than those of C57 mice [25], but this is not a driver of pathological HCM. Echocardiographic findings from Coley *et al.* [19] show no deficits in D2.WT cardiac function from 7 to 52 weeks of age. Similarly, longitudinal data of sedentary and wheel-running D2.WT mice within the current study reveal no evidence of a progressive HCM (**Figure 1-2**).

It has been asserted that D2.WT mice harbor mutations (i.e. different from the sequence expressed by C57BL/6 mice) in *Myh7* (encoding β-cardiac myosin) and myosin binding protein C (MyBP-C) that lead to HCM [19]. A close inspection of these polymorphisms reveal that this is not likely to be the case. In the case of the *Myh7* polymorphism, a new start site is established that adds five amino acids to the N-terminus without interfering with expression of the myosin. Indeed *Myh7* expression is shown in the publication by Zhao et al. [19]. Based on a large volume of work in the myosin field, tags of various size, including GFP, can be placed on the N-terminus of class II myosins (including cardiac) without any functional impact [26, 27]. Furhermore, α-cardiac myosin (*Myh6*), not β-cardiac myosin, is the predominant isoform in the adult mouse heart, with the β isoform being primarly expressed during development [28].

The situation with mouse MyBP-C is even more interesting. The publication of Zhao *et al*. [19] asserted that the normal translation start site was lost in the DBA/2J background, which would result in a truncated protein that would be pathogenic. However, if one compares mouse and human MyBP-C sequences [29], one finds that some mouse strains, including the C57BL/6, have an N-terminal extension not found in humans: <u>PGVTVLKM</u>PEPGKKPVS (the mouse-specific N-terminal extension is underlined). Interestingly the loss of the normal mouse start site will remove the mouse-specific N-terminal extension and the protein will instead start at the normal human start site (the methionine is processed off, leaving the proline as the first amino acid in the native protein). The work of Bunch *et al.* [29] demonstrated that this mouse N-terminal extension, in fact, intereferes with the interaction of the N-terminus of MyBP-C with myosin, therefore, its removal is likely beneficial. It appears that only certain strains of mice have gained a new start site (including SVE/129, C57BL/6, and FVB/N), while others, including the DBA/2J, have maintained (or regained) the human start site. Zhao *et al.* [19] also found four polymorphisms in the MyBP-C coding sequence of DBA/2J vs. C57 mice, but none of the differences have ever been linked to altered cardiac function or disease.

Herein, we domonstrate that DBA/2J mice exhibit similar perivascular connective tissue as C57 mice and healthy canines (**Figure 1a-f**). Quantification of LV collagen content reveals that DBA/2J mice have no more ECM than healthy canines and humans (**Figure 1g**). Using high frequency ultrasound, we imaged the hearts of DBA/2J mice and found no LV wall thickening with wheel running (**Figure 2a**). Mice of both activity levels had preserved ejection fraction (**Figure 2e**) and diastolic function (**Figure 2h**) throughout the 12 month long study. Our data and these considerations, thus, support the use of the DBA/2J background strain for genetic manipulations that cause cardiomopathy.

## METHODS & MATERIALS

### Animals

This study used male D2.WT (Jax# 000671) and B10.WT (Jax# 000476) mice from colonies originally obtained from Jackson Laboratory. Mice were housed 1-5 mice per cage, randomly assigned into groups, provided *ad libitum* access to food (NIH-31 Open formulation diet; Envigo #7917), water, and enrichment, and maintained on a 12-hour light/dark system. Volitional running in D2.WT mice was implemented as previously described [23]. All animal studies were approved and conducted in accordance with the University of Florida IACUC.

### Echocardiography

Transthoracic echocardiograms were performed using the Vevo 3100 pre-clinical imaging system (Fujifilm Visualsonics). Mice were anesthetized using 3% isoflurane and maintained at 1.5-2% to keep heart and respiration rates consistent among mice. Body temperature was maintained throughout imaging with the use of a heated platform and a heat lamp. Four images were acquired for each animal: B-mode parasternal long axis (LAX), B-mode short axis (SAX), M-mode SAX, and apical four-chamber view with color doppler and pulsed-wave doppler. M-mode SAX images were acquired at the level of the papillary muscle. Flow through the mitral valve was sampled at the point of highest velocity, as indicated by aliasing, with the pulsed-wave angle matching the direction of flow. Images were imported into Vevo LAB for analysis. Measurements of M-mode SAX and pulsed-wave doppler images were made from three consecutive cardiac cycles between respirations.

### Immunofluorescence and histological evaluations

Fresh-frozen OCT-embedded hearts were sectioned at 10 μm and fixed in ice-cold acetone. The sections were re-hydrated in PBS, blocked in 5% BSA-PBS at room temperature and incubated with the anti-syntrophin primary antibody (1:2000; #11425; Abcam) overnight at 4°C. Following PBS washes, sections were incubated at room temperature with a fluorescent dye-conjugated secondary antibody and coverslipped using Prolong Gold anti-fade mounting medium (ThermoFisher Scientific). Slides were visualized with a Leica DMR microscope, and images were acquired using a Leica DFC310FX camera interfaced with Leica LAS X software. Cardiomyocyte minimum ferret diameter was measured in ImageJ by investigators blinded to study groups. Picrosirius Red (PSR) staining was performed as previously described [14].

### Collagen Assay

Snap-frozen LV samples were pulverized and disrupted with a hand homogenizer in dH_2_O. A volume of 200 μL for each sample homogenate was completely desiccated by overnight incubation at 65^ο^ C to obtain the dry mass of the sample to be assayed, avoiding artifacts caused by edema. Following dry mass determination, the collagen content of the samples was measured using a colorimetric Total Collagen Assay kit (Biovision #K218) in a 96-well plate using a SpectraMax i3x multi-mode spectrophotometer (Molecular Devices). Canine samples were utilized from a previous study [5], and human samples were obtained from the National Disease Research Interchange (Philadelphia, PA).

### Statistical analysis

Statistical analysis was performed using the appropriate form of ANOVA (one-way, two-way, or repeated measures) followed by Tukey HSD post-hoc tests (α = 0.05). A P value less than 0.05 was considered significant. Data are displayed as mean ± SEM or box-and-whisker plots.

## ACKNOWLEDGMENTS

This work was funded by a Wellstone Muscular Dystrophy Cooperative Center grant (P50-AR-052646) from the NIH to HLS and DWH, a Parent Project Muscular Dystrophy grant to HLS, and a grant from the Muscular Dystrophy Association (MDA549004) to DWH. Michael Matheny, Lillian Wright, and Shailja Desai are thanked for their technical support related to this project.

## AUTHOR CONTRIBUTIONS

Study design was contributed by CCH, YL, DWH, and HLS. Experimental procedures and data acquisition were conducted by CCH, YL, and DWH. All authors were involved in data analysis, interpretation, data presentation, and manuscript writing.

## Conflict of Interested Statement

The authors have declared no conflict of interest exists.

## Data Availability

The datasets generated during and/or analyzed during the current study are available from the corresponding author on reasonable request.

## References

[1] Markati T, De Waele L, Schara-Schmidt U, Servais L. Lessons Learned from Discontinued Clinical Developments in Duchenne Muscular Dystrophy. Front Pharmacol. 2021;12:735912.

[2] Bulfield G, Siller WG, Wight PA, Moore KJ. X chromosome-linked muscular dystrophy (mdx) in the mouse. Proc Natl Acad Sci U S A. 1984;81:1189–92.

[3] Im WB, Phelps SF, Copen EH, Adams EG, Slightom JL, Chamberlain JS. Differential expression of dystrophin isoforms in strains of mdx mice with different mutations. Hum Mol Genet. 1996;5:1149–53.

[4] de Lucia C, Wallner M, Eaton DM, Zhao H, Houser SR, Koch WJ. Echocardiographic Strain Analysis for the Early Detection of Left Ventricular Systolic/Diastolic Dysfunction and Dyssynchrony in a Mouse Model of Physiological Aging. J Gerontol A Biol Sci Med Sci. 2019;74:455–61.

[5] Hammers DW, Sleeper MM, Forbes SC, Shima A, Walter GA, Sweeney HL. Tadalafil Treatment Delays the Onset of Cardiomyopathy in Dystrophin-Deficient Hearts. J Am Heart Assoc. 2016;5.

[6] Yasuda S, Townsend D, Michele DE, Favre EG, Day SM, Metzger JM. Dystrophic heart failure blocked by membrane sealant poloxamer. Nature. 2005;436:1025–9.

[7] Townsend D, Blankinship MJ, Allen JM, Gregorevic P, Chamberlain JS, Metzger JM. Systemic administration of micro-dystrophin restores cardiac geometry and prevents dobutamine-induced cardiac pump failure. Mol Ther. 2007;15:1086–92.

[8] Parvatiyar MS, Marshall JL, Nguyen RT, Jordan MC, Richardson VA, Roos KP, et al. Sarcospan Regulates Cardiac Isoproterenol Response and Prevents Duchenne Muscular Dystrophy-Associated Cardiomyopathy. J Am Heart Assoc. 2015;4.

[9] Grady RM, Teng H, Nichol MC, Cunningham JC, Wilkinson RS, Sanes JR. Skeletal and cardiac myopathies in mice lacking utrophin and dystrophin: a model for Duchenne muscular dystrophy. Cell. 1997;90:729–38.

[10] Sacco A, Mourkioti F, Tran R, Choi J, Llewellyn M, Kraft P, et al. Short telomeres and stem cell exhaustion model Duchenne muscular dystrophy in mdx/mTR mice. Cell. 2010;143:1059–71.

[11] Mourkioti F, Kustan J, Kraft P, Day JW, Zhao MM, Kost-Alimova M, et al. Role of telomere dysfunction in cardiac failure in Duchenne muscular dystrophy. Nat Cell Biol. 2013;15:895–904.

[12] Fukada S, Morikawa D, Yamamoto Y, Yoshida T, Sumie N, Yamaguchi M, et al. Genetic background affects properties of satellite cells and mdx phenotypes. Am J Pathol. 2010;176:2414–24.

[13] Coley WD, Bogdanik L, Vila MC, Yu Q, Van Der Meulen JH, Rayavarapu S, et al. Effect of genetic background on the dystrophic phenotype in mdx mice. Hum Mol Genet. 2016;25:130–45.

[14] Hammers DW, Hart CC, Matheny MK, Wright LA, Armellini M, Barton ER, et al. The D2.mdx mouse as a preclinical model of the skeletal muscle pathology associated with Duchenne muscular dystrophy. Sci Rep. 2020;10:14070.

[15] Rodrigues M, Echigoya Y, Maruyama R, Lim KR, Fukada SI, Yokota T. Impaired regenerative capacity and lower revertant fibre expansion in dystrophin-deficient mdx muscles on DBA/2 background. Sci Rep. 2016;6:38371.

[16] Mazala DA, Novak JS, Hogarth MW, Nearing M, Adusumalli P, Tully CB, et al. TGF-beta-driven muscle degeneration and failed regeneration underlie disease onset in a DMD mouse model. JCI Insight. 2020;5.

[17] Hakim CH, Wasala NB, Pan X, Kodippili K, Yue Y, Zhang K, et al. A Five-Repeat Micro-Dystrophin Gene Ameliorated Dystrophic Phenotype in the Severe DBA/2J-mdx Model of Duchenne Muscular Dystrophy. Mol Ther Methods Clin Dev. 2017;6:216–30.

[18] Nabors CE, Ball CR. Spontaneous calcification in hearts of DBA mice. Anat Rec. 1969;164:153–61.

[19] Zhao W, Zhao T, Chen Y, Zhao F, Gu Q, Williams RW, et al. A Murine Hypertrophic Cardiomyopathy Model: The DBA/2J Strain. PLoS One. 2015;10:e0133132.

[20] Heydemann A, Ceco E, Lim JE, Hadhazy M, Ryder P, Moran JL, et al. Latent TGF-beta-binding protein 4 modifies muscular dystrophy in mice. J Clin Invest. 2009;119:3703–12.

[21] Ceco E, Bogdanovich S, Gardner B, Miller T, DeJesus A, Earley JU, et al. Targeting latent TGFbeta release in muscular dystrophy. Sci Transl Med. 2014;6:259ra144.

[22] Maron BJ, Ackerman MJ, Nishimura RA, Pyeritz RE, Towbin JA, Udelson JE. Task Force 4: HCM and other cardiomyopathies, mitral valve prolapse, myocarditis, and Marfan syndrome. J Am Coll Cardiol. 2005;45:1340–5.

[23] Hammers DW, Sleeper MM, Forbes SC, Coker CC, Jirousek MR, Zimmer M, et al. Disease-modifying effects of orally bioavailable NF-kappaB inhibitors in dystrophin-deficient muscle. JCI Insight. 2016;1:e90341.

[24] Marian AJ. Molecular Genetic Basis of Hypertrophic Cardiomyopathy. Circ Res. 2021;128:1533–53.

[25] Rosenkranz S. TGF-beta1 and angiotensin networking in cardiac remodeling. Cardiovasc Res. 2004;63:423–32.

[26] Wolny M, Colegrave M, Colman L, White E, Knight PJ, Peckham M. Cardiomyopathy mutations in the tail of beta-cardiac myosin modify the coiled-coil structure and affect integration into thick filaments in muscle sarcomeres in adult cardiomyocytes. J Biol Chem. 2013;288:31952–62.

[27] Kengyel A, Wolf WA, Chisholm RL, Sellers JR. Nonmuscle myosin IIA with a GFP fused to the N-terminus of the regulatory light chain is regulated normally. J Muscle Res Cell Motil. 2010;31:163–70.

[28] Ng WA, Grupp IL, Subramaniam A, Robbins J. Cardiac myosin heavy chain mRNA expression and myocardial function in the mouse heart. Circ Res. 1991;68:1742–50.

[29] Bunch TA, Lepak VC, Kanassatega RS, Colson BA. N-terminal extension in cardiac myosin-binding protein C regulates myofilament binding. J Mol Cell Cardiol. 2018;125:140–8.

